# Microplastics drive both linear and threshold-type shifts in soil multifunctionality along concentration gradients

**DOI:** 10.64898/2026.01.31.701563

**Authors:** Tamara Meizoso-Regueira, Marina Dacal, Emma Ring, Matthias C. Rillig

## Abstract

Microplastics are increasingly recognized as emerging contaminants in terrestrial ecosystems, yet their mechanistic impacts on soil multifunctionality remain poorly understood. Here, we evaluated the influence of two microplastic polymers, polyethylene terephthalate and polypropylene, on soil functioning by subjecting soils to a gradient of concentrations of these microplastics, and measuring six variables representing soil physical, chemical, and biological functions. A statistical framework combining multi-model inference with threshold detection and machine learning was implemented in this study to identify the main pathways of soil multifunctional change. Most significant responses followed nonlinear trends and threshold shifts, primarily in physical properties, indicating that microplastic stress first impacts soil structure before cascading to chemical and biological processes. We identified two system-level thresholds at 0.3% PP and 0.55% PET w/w; while random forest highlighted water-stable aggregates as the dominant predictor of overall soil multifunctionality. Our findings provide new quantitative evidence of complex soil multifunctionality responses to microplastic pollution. Most importantly, physical deterioration emerged as an early-warning signal of microplastic disturbance, thereby advancing our understanding of microplastic pollution on soil systems.

## 1. Introduction

The plastic industry has revolutionized our society since the Second World War, shaping how we live and interact with our environment^1^. The versatility, low cost, and durability of plastic have enabled an enormous progress across numerous sectors, including technology, medicine, textiles, and agriculture^2–4^. However, their extensive production and widespread use have led to plastic pollution becoming a global environmental challenge. Once plastics are released and accumulated in ecosystems, they gradually fragment and degrade through physical and biochemical processes, generating particles smaller than 5 mm, known as microplastics^5,6^. These particles are often classified as secondary microplastics, while primary microplastics are those intentionally manufactured at microscopic sizes for industrial or cosmetic applications^3^.

One of the most important challenges of plastic pollution is its environmental persistence. Most polymers require hundreds to thousands of years to degrade completely, leading to a continuous accumulation of plastics over time in both aquatic and terrestrial ecosystems^7–9^. Although past research efforts have focused predominantly on marine and freshwater ecosystems, soils have increasingly been recognized as major sinks for microplastics^10,11^. In terrestrial ecosystems, microplastics enter through multiple pathways, including agricultural practices like compost application, plastic mulching or irrigation with contaminated water, as well as poor waste management or atmospheric deposition^12,13^. Once in the soil, microplastics persist, migrate, and interact with soil components in complex ways^12,14^.

Previous scientific evidence indicates that microplastics influence soil physical, chemical, and biological properties. These impacts include changes in soil aggregation and pore structure, as microplastics may interfere with particle binding and pore connectivity, altering water infiltration and retention^15–17^. Microplastics have also been associated with reductions in enzymatic activity and soil respiration, affecting microbial communities and carbon cycling processes^18–21^, since they can modify nutrient availability, as well as introduce additives that disturb microbial functioning. In addition, microplastics can also impact plant growth and performance through changes in nutrient availability, reduced water uptake, or affecting rhizosphere biota^22,23^. Importantly, these responses are not always consistent across studies, as they depend on polymer type, particle shape, size, and surface chemistry, and soil characteristics, resulting in both inhibitory and, occasionally, stimulatory responses^15,16,24^.

Despite these advances, the understanding of microplastic dynamics in soil remains limited. Although some studies have begun to explore the complexity of microplastic impacts, including differences among polymer types, particle sizes and shapes^24,25^, as well as interactions with other stressors such as drought and heavy metals^26,27^, existing evidence remains incomplete and does not yet provide a coherent mechanistic explanation. More recently, it has been suggested that microplastics, similarly to other global change factors, may induce nonlinear or threshold responses in soil properties, where abrupt shifts occur once certain concentration levels are exceeded^28,29^. Such behaviour has been observed in other ecological contexts; for instance, Scheffer et al.^30^ demonstrated that biodiversity can enhance the resistance of ecosystem productivity to climate extremes. Similarly, soils may buffer low levels of disturbance through compensatory processes, such as microbial or structural reorganization, under which soil functioning remains relatively stable until a critical tipping point is surpassed. Once this buffering capacity is exceeded, potential system-level shifts, sometimes irreversible^31^, may occur. In this context, microplastics may even induce hormetic patterns, stimulating positive responses at low concentration before negative effects emerge^32,33^. However, threshold-type soil responses to microplastics have rarely been addressed before, despite its importance for distinguishing gradual responses from abrupt shifts that may signal a critical loss of soil functionality and resilience.

Most studies aiming to identify microplastic thresholds in terrestrial ecosystems typically examine only a narrow concentration range^18,29^, focusing on a single polymer type or shape, or evaluate responses at only one timepoint. In addition, linear or monotonic dose-response models are commonly applied, which limits the ability to detect abrupt or nonlinear changes, possibly hiding potential tipping points. Compared to aquatic ecosystems, threshold-oriented experiments in terrestrial environments remain scarce, leaving concentration-response patterns in soils poorly characterized. As a result, it is still unclear whether soil responses to increasing microplastic contamination tend to be gradual or whether they exhibit abrupt shifts indicative of ecological thresholds^28,34^, and major knowledge gaps still remain. Addressing this question is essential for incorporating microplastics into risk assessment frameworks.

Beyond identifying critical concentrations for individual properties, it is important to evaluate multiple soil responses as interdependent functions to uncover integrated patterns at the ecosystem level^18,27^, where physical structure, microbial activity, and biogeochemical processes interact in complex ways^18,35,36^. Nevertheless, multifunctionality analyses under controlled microplastic concentration gradients are extremely limited, and no study has systematically tested whether different soil functions exhibit coordinated thresholds. Together, these knowledge gaps highlight the need for experimental designs that combine broad concentration gradients with flexible, nonlinear modelling capable of detecting both gradual and abrupt responses across several soil functions and soil multifunctionality.

In this study, we aimed to determine whether microplastic contamination produces gradual or abrupt shifts in key soil functions and in overall soil multifunctionality. We performed an incubation experiment using two microplastic types, PET and PP, at a concentration gradient including 20 levels, from 0 to 0.95% w/w, equally-spaced with 0.05% increments. We measured different physical, chemical and biological soil functions such as soil respiration, two enzymatic activities, pH, water-stable aggregates and mean weight diameter of aggregates, to perform a multifaceted characterization of soil responses to this kind of disturbance. We further calculated soil multifunctionality to account for the interaction between the responses of different soil functions to microplastic pollution. We conducted the experiment at three incubation periods (two, four, and six weeks) to evaluate whether soil responses to such disturbance vary with time. More specifically, we aimed to: (i) assess whether individual soil functions and multifunctionality follow linear, nonlinear, or threshold-like responses along the microplastic gradient; (ii) determine whether these patterns differ between polymers and incubation times; and (iii) interpret how response shapes develop for the microplastic concentration gradient and understand the underlying mechanisms.

## 2. Materials and methods

### 2.1. Soil and microplastic particles

The test soil was collected down to 50 cm depth from a grassland site (52°47 0N, 13°30 0E) belonging to the Institute of Biology, Free University of Berlin. It is a sandy loam soil classified as an Albic Luvisol according to the USDA soil taxonomy^37^. This site is located in a temperate oceanic climate region, characterized by a mean annual temperature of approximately 10 °C and an average annual precipitation of about 550–600 mm^38^, and the vegetation coverage is mostly grass. Fresh soil was dried in a 60°C oven for one week, sieved through a 2-mm mesh sieve and homogenized. Soil water holding capacity (WHC) was 38.5% and pH 5.8.

For this study, we selected two microplastic types, polyethylene terephthalate (PET) and polypropylene (PP), because they are among the most commonly produced and environmentally widespread polymers globally, frequently detected in both agricultural and natural soils. PET was used as powder particles (BAM code 5.3-15116) with a mean size of 211.5±131.6 µm, while PP was utilized in granular fragments (BAM code 5.3-15114) with an average size of 474.7±243.3 µm. Both microplastics were obtained from the *Bundesanstalt für Materialforschung und -prüfung* (BAM, Berlin, Germany). To minimize microbial contamination, microplastics were sterilized by rinsing with 70% ethanol for 1 minute and heating in an oven at 101°C for 48 h.

### 2.2. Experimental set-up

For the experiment, 30 g of test soil were mixed with either PET powder particles or PP granular fragments at 20 different concentration levels from 0 to 0.95% (w/w) with 0.05% increments, with the first level with no microplastic addition (0% w/w) serving as the control group. More precisely, microplastic concentrations were mixed with the soil by stirring with a metal spatula for three minutes before transferring the mixture to 50-ml polypropylene centrifuge tubes (Corning Mini Bioreactor 431720, Corning Incorporated). These tubes had lids with four vents and a hydrophobic membrane to allow gas exchange while preventing moisture loss and contamination. Control samples were stirred in the same way to ensure equivalent disturbance and minimize potential confounding effects associated with this process. We set up four replicates per treatment combination, resulting in 160 samples (20 concentration levels × two polymer types × four replicates). We replicated this experimental design for three different incubation periods (i.e., two, four, and six weeks) to assess whether the impacts of microplastics on soil biotic and abiotic properties varied with time, giving a total of 480 samples. All samples were incubated at 25°C, 60% water holding capacity (WHC) and in dark conditions.

Throughout the incubation period, all tubes were randomly distributed in the climate chamber. Harvest and analyses were also performed following this random set-up. Soil moisture was maintained at 60% WHC throughout the experiment by compensating for evaporation losses weekly, adding distilled water to each tube based on its weight loss. At each harvest time, soil respiration was measured using the complete fresh sample. Then, a portion of the sample was kept at 4°C for enzymatic activity analyses, and the rest of the sample was air-dried for a month for physicochemical analyses such as pH, water-stable aggregates and mean weight diameter measurements.

### 2.3. Soil biotic and abiotic functions

#### Soil respiration

Soil CO₂ efflux was quantified as a proxy of soil microbial respiration. These measurements were conducted at the end of each incubation period (two, four and six weeks) on the intact fresh soils before destructive sampling. More precisely, the original caps of the experimental tubes were replaced with modified rubber-septum caps (VWR, Germany; 548-3369) to seal the tubes. The headspace of the tubes was flushed for 5 min with CO₂-free air to eliminate accumulated CO_2_ and ensure that measurements reflected microbial basal respiration. Then, samples were incubated at 30°C for four hours in dark conditions. After incubation, 1 mL gas was sampled from each tube headspace to determine CO_2_ concentration using a Li-COR 6400XT infrared gas analyzer (Li-COR, USA).

#### Enzymatic activities

β-D-glucosidase and cellobiohydrolase were assessed using 5 g of soil, following a high-throughput microplate protocol defined by Jackson et al.^39^. Briefly, each sample was homogenized in 8 mL of 50 mM acetate buffer (pH 5.0–5.4) and vortexed. Then, 150 µL aliquots of the soil slurry were distributed into several wells of a 96-deep-well microplate, with some receiving substrate and others serving as controls containing only acetate buffer. Two substrates were used: 5 mM 4-p-nitrophenyl-β-D-glucopyranoside for β-D-glucosidase and 2 mM 4-p-nitrophenyl-β-D-cellobioside for cellobiohydrolase (Sigma, Germany; N7006, N5759), each added as 150 µL aliquots per well. Plates were incubated at 25°C under dark conditions for 2 h and 4 h for β-D-glucosidase and cellobiohydrolase, respectively. After incubation, they were centrifuged at 3000 rpm for 5 min, and 100 µL of the supernatant was transferred to a new plate containing 200 μL of 0.05 M NaOH in each well. Absorbance was measured at 410 nm using a Benchmark Plus microplate reader (Bio-Rad, USA).

#### Soil pH

This property was determined following the method of Carter and Gregorich^40^. Air-dried soil was mixed with a 0.01 M CaCl_2_ at a 1:2.5 ratio in 50 mL tubes and vortexed for 30 min. The tubes were then centrifuged for 10 min at 4600 rpm at room temperature to ensure complete sedimentation of soil particles, and 20 mL of the supernatant was transferred to new centrifuge tubes (Sarstedt AG & Co. KG, Nümbrecht, Germany; item no. 62.548.004) and used for pH determination with a Knick 766 pH meter (Knick, Germany).

#### Water-stable aggregates (WSA)

Aggregate stability was quantified based on Kemper and Rosenau^41^. Air-dried soil samples are weighed to 4 g from the whole sample, then pre-wetted with deionized water, placed on 250 µm sieves, and oscillated vertically in a wet-sieving apparatus for 3 min. After sieving, the material was dried overnight (60 °C) and weighed to obtain the “dry matter” fraction. Samples were then washed through a smaller sieve to remove sand and organic debris, dried again, and re-weighted as the “coarse matter” fraction. Then the WSA (%) was calculated as follows: WSA=[(dry matter – coarse matter)/(4 g – coarse matter)]*100.

#### Mean weight diameter (MWD)

Aggregate size distribution was also determined on the air-dried soil following the dry-sieving protocol of Kemper and Rosenau^41^. Soil samples were passed through a stack of sieves with decreasing mesh sizes (2 mm, 1 mm, 250 µm, 53 µm). The mass retained on each sieve was weighed after shaking the sieve stack horizontally until complete separation. Each fraction was estimated based on the corresponding mesh size (i.e. 2 mm weight/total sample weight; 1 mm weight/total sample weight, etc.). The mean diameter for each fraction was defined as the midpoint between the two sieve sizes, and mean weight diameter was calculated as the weighted mean of the fractions using the sieve size midpoint as the weighting factor, using the following formular: MWD=((fraction1*3)+(fraction2*1.5)+(fraction3*0.625)+(fraction4*0.1515)+(fraction5*0.0265)).

### 2.4. Multifunctionality index

We quantified overall ecosystem performance using a soil multifunctionality index (MFI), which is also defined as the ecosystem ability to sustain multiple functions simultaneously, following a standardized approach already consolidated in previous studies^35,42,43^. This approach has previously been applied to microplastic–soil studies by Cheng et al.^18^. For each timepoint, all soil properties considered in this study were first standardized using Z-scores to remove scale differences among functions. After standardization, the MFI was calculated for each sample as the arithmetic mean of the six standardized functions. This approach provides an integrated measure of soil functioning that accounts for physical, chemical, and biological processes simultaneously (with equal weighting), enabling comparison of microplastic effects on the soil system as a whole rather than on isolated properties.

### 2.5. Statistical analysis

Summary statistics were examined to detect skewness, kurtosis, and heteroscedasticity patterns. Outliers were identified using the 1.5×IQR rule, and their influence was evaluated through a complementary analysis of robustness as a sensitivity check. OLS were applied on the IQR-cleaned dataset, while we refitted the models on the raw dataset using Huber M-estimation, which downweighs the outliers without removing them. We then compared significance and effect direction against the OLS fits (without outliers). As outliers did not significantly affect model selection outcomes in most cases, they were excluded from the final models to simplify interpretation.

To address whether soil functions respond to a gradient of microplastic concentrations in a gradual or abrupt way, we applied a multi-model selection workflow for threshold detection based on Berdugo et al^34^. We first assessed linear vs. nonlinear relationships by fitting a linear model *versus* a quadratic polynomial and a Generalized Additive Model (GAM). Unlike Berdugo et al.^34^, our model selection based on AIC comparison was complemented with a conservative parsimony-based approach: if two models differed by less than two AIC units (ΔAIC ≥ 2), we chose the simpler model as the most parsimonious explanation, in line with Burnham and Anderson^44^.

When nonlinearity was supported against the linear model by this AIC criteria, we compared the best nonlinear fit against a family of changepoint (piecewise) candidates: hinge (constant slope before or after the breakpoint), segmented (slope change on both sides of the breakpoint), step (sudden jumps in a response variable without local slope change) and stegmented (sudden shift in both intercept and slope)^45^. A piecewise model and its estimated breakpoint were accepted following the same approach (when its AIC was at least two units lower than the nonlinear AIC (ΔAIC ≥ 2). The breakpoint location (ψ) was estimated by selecting the ψ candidate that minimized AIC among all evaluated piecewise models. Breakpoint uncertainty was assessed using profile-AIC uncertainty intervals^46^, representing the range of ψ values that fit the data nearly as well as the best estimate (ΔAIC ≤ 2). These intervals reflect model-fit uncertainty rather than classical sampling-based confidence intervals; bootstrap-based confidence intervals were explored but were found to be unstable in our data.

To ensure that only meaningful responses were interpreted as microplastic-driven, model significance was evaluated through harmonized criteria depending on model type: F-tests comparing each model against its null (for linear and quadratic), or smooth and slope/step term significance (for GAMs and piecewise models, respectively). The aim of this step was not to filter out non-significant results, but to distinguish between two statistically different questions: (i) which model best describes the shape of the data according to AIC, and (ii) whether that shape can genuinely be attributed to the microplastic concentration gradient. AIC identifies the model with the best relative fit, but does not guarantee that the corresponding model shape can be attributed to the microplastic concentration gradient. Flexible models, such as GAMs or piecewise regressions, can win by AIC simply because they are more flexible and adapt better to random noise, even when the data show no meaningful trend^44^, potentially leading to overfitting or type-I error inflation.

Therefore, models were considered significant (i.e. the microplastic concentration is the factor driving the change) when either the F-test p-value was < 0.05, or the smooth/slope term p-value was < 0.05 in those cases where F-test is not meaningful. For linear and quadratic models, we initially screened significance based on the slope term. However, model selection decisions were ultimately based on the overall F-test, which provides a stricter and more conservative criterion for testing whether microplastic concentration had a significant effect on the response. The F-test, or an equivalent global test, could not be directly interpreted from GAMs and piecewise regressions because these models are based on nonlinear smooth or discontinuous terms that lack a single overall slope or parametric degree of freedom for comparison (equivalent global tests are less stable and comparable). In those cases, the concentration effect p-value corresponds to the significance of the smooth term for GAMs; for piecewise models, it corresponds to the significance of at least one slope or step parameter, indicating whether the response changes significantly before or after the estimated breakpoint.

To evaluate system-level responses to microplastics, we also estimated overall ecosystem performance with a soil multifunctionality index (MFI) following the approach described above and widely adopted in soil multifunctionality literature^18,35,36,42,43^. The same multi-model selection framework applied to individual properties was followed for MFI. Besides, to identify which soil functions most strongly influence multifunctionality, random forest analysis (1,000 trees) was performed using standardized functions as predictors to identify the most influential variable contributing to MFI. To ensure robustness, forests were repeated several times (1000 runs) using different random seeds, and predictor importance was summarized by the mean permutation importance across runs. Statistical significance of top predictors was evaluated by empirical p-values based on the distribution of permutation importance across repeated forests, and 95% confidence intervals were obtained from the corresponding empirical quantiles. Model performance was evaluated using out-of-bag R².

Finally, to test mechanistic pathways linking microplastic concentration to multifunctionality based on the predicted most-influential variables, potential mechanistic links were explored using bootstrapped mediation analysis^47^. This approach is adapted from behavioural causal-inference theory, and evaluates whether changes in key soil properties may act as intermediate pathways linking microplastic concentration to multifunctionality. More precisely, microplastic concentration was treated as the predictor, the predicted influential variables as mediators, and MFI as the response variable. Indirect effects (a·b) were estimated using ordinary least squares (OLS) models with percentile bootstrap confidence intervals (B ∼ 2000), empirical two-sided p-values, and model R² reported for each path. In the case of significant mediation, as a robustness check, the analysis was repeated excluding the top variable from the MFI computation, to verify whether that variable acted as a mediator beyond its direct contribution to the index.

All analyses were conducted in R (version 4.5.2). Data handling and pre-processing was conducted using the packages *dplyr*^48^, *tidyr*^49^, *tibble*^50^, *stringr*^51^, and *glue*^52^. Model fitting was performed with the package *mgcv*^53^ for generalized additive models, the *lm* function from the *base* package^54^ for linear and polynomial models, and custom AIC-based workflows for piecewise regressions. Random forests were fitted using the package *ranger*^55^. Statistical testing and model summarization were performed with the packages *broom*^56^, *MASS*^57^, and *sandwich*^58^. Finally, for visualization and figure composition, the packages *ggplot2*^59^, *cowplot*^60^, and *patchwork*^61^ were used.

## 3. Results

### 3.1. Gradual and abrupt responses of soil functions along the concentration gradient

Regarding the dose-response relationships across individual functions, the most frequent winner model by AIC comparison was linear (Table S1, S2 and S3) at first; however, after applying the significance criteria previously described, the remaining meaningful models were mostly piecewise and nonlinear, indicating that abrupt responses were actually more common than suggested by AIC alone. The significance criteria to collect only meaningful models were applied not to favour significant results, but to distinguish between relative model fit and evidence for a genuine concentration effect. Because flexible models can outperform simpler ones by AIC even when patterns arise from noise, we additionally required a significant concentration term (F-test and/or slope/smooth term p-value < 0.05) to avoid interpreting overfitted models as microplastic effects. Non-significant models for individual properties (Fig. S1, S2, S3 and S4), as well as for MFI (Fig. S5), are still reported in the Supplementary Materials.

This way, we observed that the PET concentration gradient produced mostly gradual changes in MWD (Fig. 1a, 1b), while PP consistently induced abrupt shifts (Fig. 1d, 1e) in aggregate size with increasing concentrations of this microplastic. Three additional variables showed significant responses to the microplastic gradient: a step-like decrease in CB under PP (Fig. 1f), a segmented shift in pH under PET (Fig. 1g), and a linear decline in WSA under PP (Fig. 1h). Most significant responses in soil properties occurred after four to six weeks of incubation. Specifically, most abrupt responses emerged after six weeks of incubation, with thresholds emerging at low, medium, and high concentration levels depending on the property, polymer and timepoint combination. Most of these abrupt relationships showed step-like thresholds, reflecting sudden transitions between low and high functional states before or after the breakpoint. Nevertheless, profile-AIC uncertainty intervals expand to the full search range (± 0.20) for individual functions, indicating weak identifiability of the exact breakpoint (Table 1). Thus, while abrupt shifts were supported, the precise breakpoint location remains uncertain.

**Figure 1.**
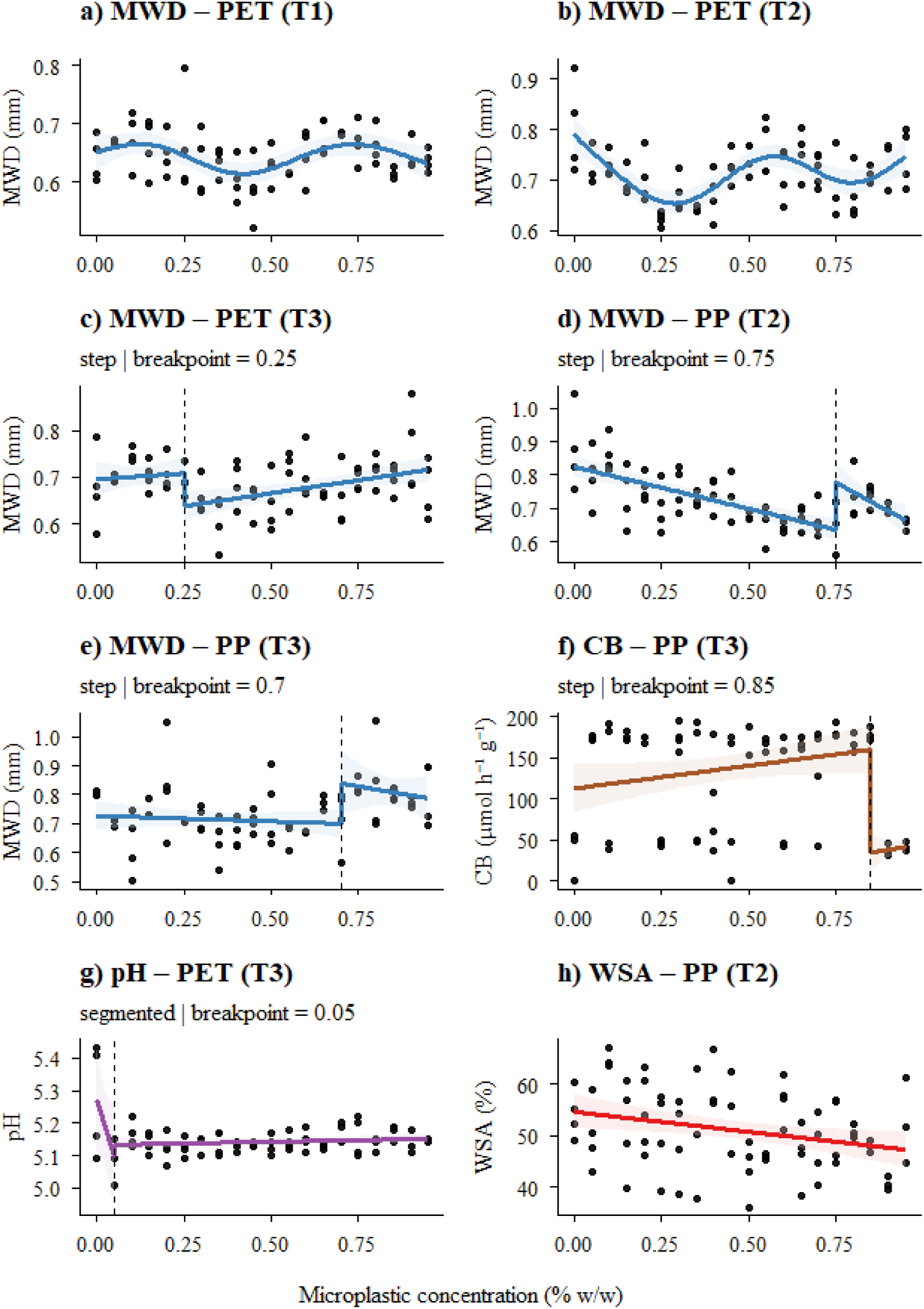
Individual functions winner-models by AIC comparison that passed the significance criteria. PET: polyethylene terephthalate, PP: polypropylene. T1, T2 and T3 correspond to 2, 4, and 6 weeks of incubation, respectively. Piecewise (panels c-g) and GAM (panels a-b) models presented a significant p-value of the smooth and slope/step term, respectively; while the linear model (panel h) had a significant p-value from the model F-test. Discontinuous lines show estimated breakpoints.

**Table 1.**
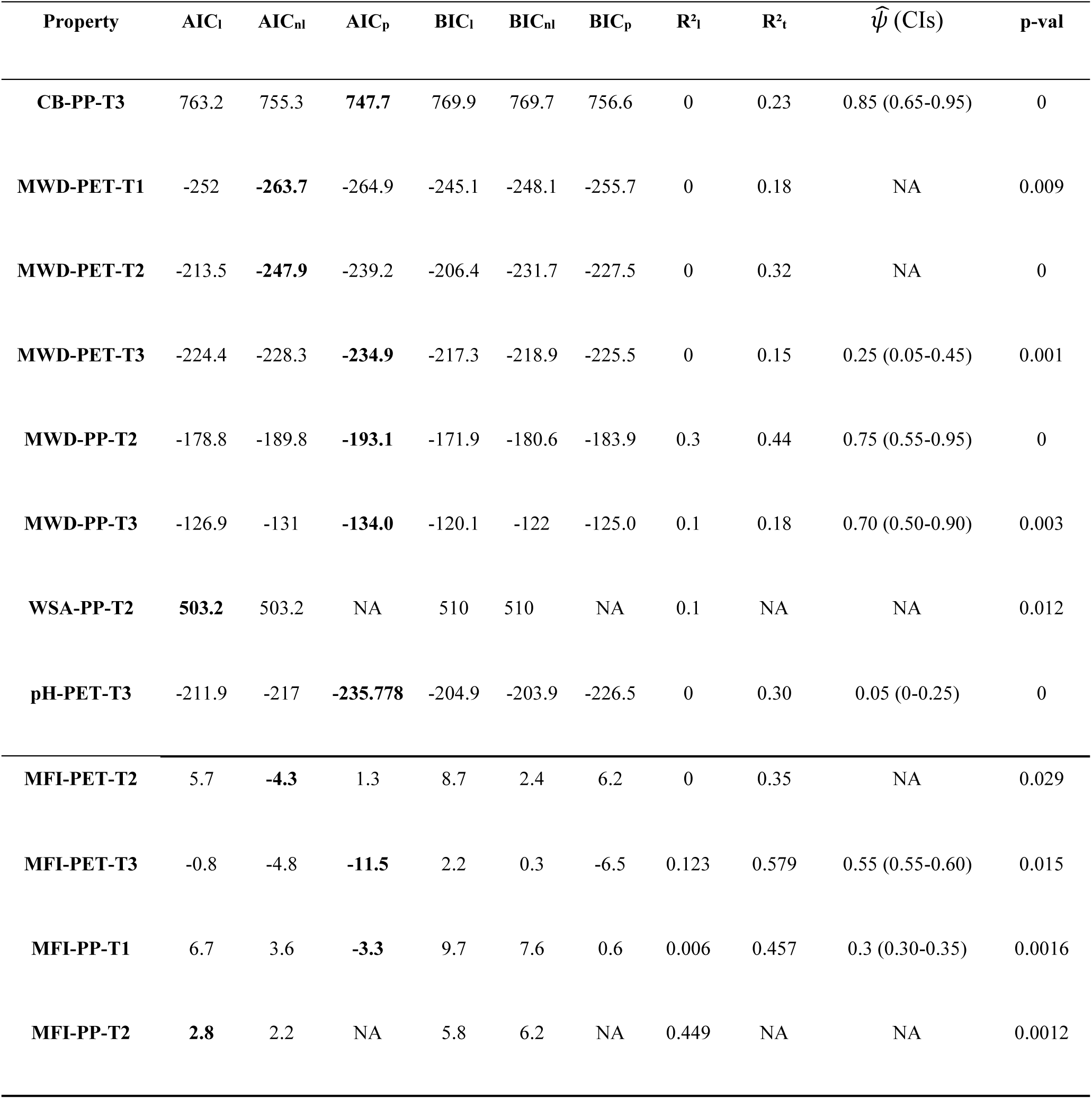
Goodness-of-fit metrics to evaluate and compare model fit for each combination of property x polymer x timepoint, as well as MFI x polymer x timepoint, that passed the significance criteria, including AIC, BIC and R². Highlighted in bold are the winner models following our ΔAIC ≥ 2 criteria, prioritizing parsimony (l=linear, nl=nonlinear, p=piecewise). The term p-val corresponds to the overall F-test for linear and quadratic models, or to the significance of the smooth and slope/step term for GAM and piecewise models, respectively. We report the estimated breakpoints *ψ̂* along with profile-AIC uncertainty intervals. When linear and non-linear models presented the best fit, there was no threshold to be reported.

In Table 1, for both MWD-PET-T1 and MWD-PET-T2, GAM metrics show the best fit, while linear responses were only retained in one case (WSA-PP-T2), indicating a clear monotonic concentration effect without an abrupt transition. For respiration and β-D-glucosidase, responses were mostly gradual and linear; however, linear models did not explain significantly more variance than a null model (intercept-only, no MP concentration) by F-test, so their goodness-of-fit metrics are reserved for the supplementary materials (Table S1). Taken together, soil functions displayed a mixture of gradual nonlinear and threshold responses, with physical indicators (MWD, WSA) showing the strongest evidence for abrupt transitions.

### 3.2. Direction and magnitude of microplastic impacts on soil functions across polymers

Beyond model fit, the direction and magnitude of microplastic effects on individual functions also exhibited clear differences across polymers. This way, PP consistently reduced soil structural integrity, producing declines in WSA and shifts in MWD across the microplastic gradient, with MWD dropping sharply when concentrations exceeded ∼0.7 % w/w. PET also led to a step breakpoint on MWD response (ψ = 0.25% w/w) after six weeks of incubation (Fig. 1c, Table 1), producing a modest increase in MWD after the estimated threshold, consistent with a possible reorganization of soil structure.

After six weeks of incubation, PP produced an initial plateau followed by a sharp decline at high concentrations (0.85% w/w) of CB, indicating inhibition of cellobiohydrolase. Besides, PET also induced a segmented pH response after six weeks of incubation, with a breakpoint at very low doses (ψ = 0.05 % w/w; Fig. 1g, Table 1), showing a microplastic-induced acidification at very low doses followed by a stabilization phase. WSA in soils exposed to PP after four weeks presented a significant linear decrease (Fig. 1h), demonstrating that even at low doses, PP gradually reduces the proportion of water-stable aggregates, reinforcing that physical soil degradation precedes biochemical disturbance. Overall, soil structural properties were the earliest and most indicative of microplastic stress.

### 3.3. Direction and magnitude of microplastic impacts on soil functions over time

Microplastic impacts on individual properties also differed across timepoints, intensifying with incubation time. After two weeks of microplastic exposure, dose–response patterns were weak or emerging. By week four, significant declines in MWD and WSA appeared under PP (Fig. 1b, 1d, 1h). After six weeks of incubation, most abrupt transitions, such as MWD under PET and PP (Fig. 1c, 1e), CB under PP (Fig. 1f), and pH under PET (Fig. 1g), were expressed. Combining both polymer and time-dependent effects aforementioned, we can conclude that PP seems to produce stronger and earlier declines across variables, whereas abrupt PET effects tended to appear later and with a lower magnitude.

### 3.4. Multifunctionality responses and main drivers of multifunctionality decline

Following the same multi-model comparison workflow, the multifunctionality index (MFI) presented two winner piecewise models that met the AIC and significance criteria: PET after six weeks of incubation displayed a stegmented relationship with ψ = 0.55% w/w (Fig. 2b), while PP after two weeks showed a step threshold at ψ = 0.30% w/w (Fig. 2c). In Table 1, the narrower uncertainty intervals for MFI show stronger breakpoint identifiability than for individual functions. These abrupt shifts indicate that relatively low microplastic doses can trigger coordinated change across multiple functions, and this system-level response can emerge after different exposure times depending on the microplastic type (Fig. 2b, 2c), such as quite early for PP, or only after cumulative exposure for PET. Additionally, after four weeks, MFI showed a significant linear decline (Fig. 2d) under PP exposure, suggesting a gradual loss of multifunctionality under intermediate PP doses; differently, MFI showed a nonlinear pattern under PET (Fig. 2a). Across the four significant MFI responses, the strongest system-level decline occurred under PP after four weeks (Fig. 2d), showing a ∼0.6 SD decrease across the gradient, considered moderate to strong according to Sawilowsky, 2009^62^.

**Figure 2.**
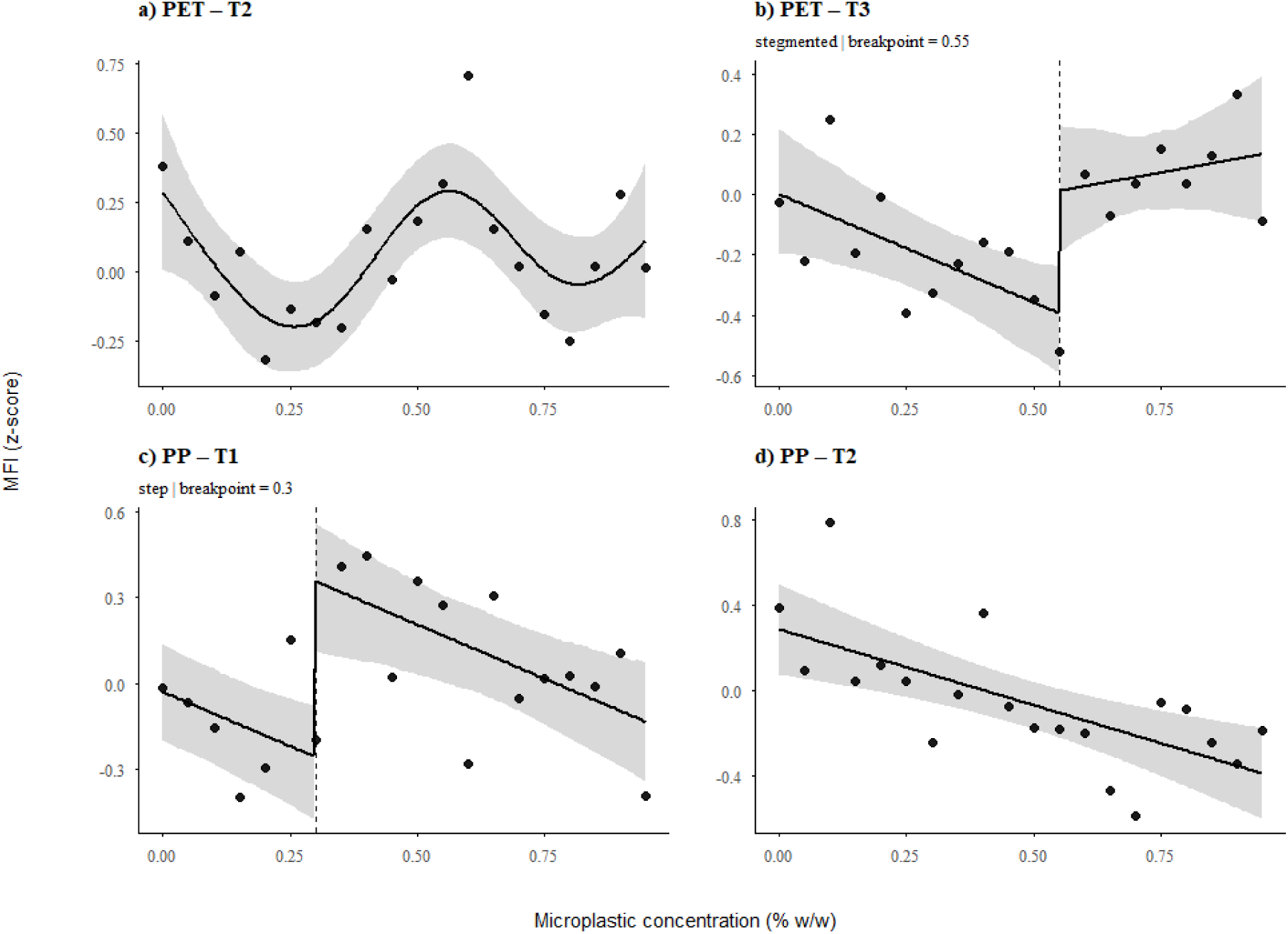
Soil multifunctionality index (MFI) winner models by AIC comparison that passed the significance criteria. PET: polyethylene terephthalate, PP: polypropylene. T1, T2 and T3 correspond to 2, 4, and 6 weeks of incubation, respectively. Panel a) MFI corresponding to PET-Timepoint 2 (GAM); b) PET-Timepoint 3 (estimated stegmented threshold around 0.55% w/w); c) PP-Timepoint 1 (estimated step-like threshold at 0.3% w/w); and d) PP-Timepoint 2 (with a significant linear trend by F-test). Negative values indicate below-average multifunctionality across soil functions (z-score units)^70^.

Random forests explained a moderate fraction of MFI variation for PP (R^2^ = 0.45-0.53; Fig. 3c, 3d), while PET exhibited weaker predictive performance across timepoints (R^2^ = 0.22-0.24; Fig. 3a, 3b). This polymer asymmetry follows the patterns observed for individual functions, where PP consistently degraded structural variables more strongly and earlier, which translated into clear system-level declines. PET, in contrast, required longer microplastic exposure before producing detectable multifunctionality disruption. In this case, respiration was identified as the most important predictive variable after four weeks, MWD ranking second after both two and four weeks (Fig. 3a, 3b). Under PP exposure, variable importance profiles were dominated by WSA as the strongest predictor, followed by MWD after four weeks (Fig. 3d), indicating the overall soil structure control on ecosystem multifunctionality. This suggests that, once microplastics first compromise aggregate stability by reducing water-stable aggregates or fragmenting aggregates, biogeochemical functions tend to decline accordingly, thereby compromising the ability of ecosystems to sustain multiple functions simultaneously.

**Figure 3.**
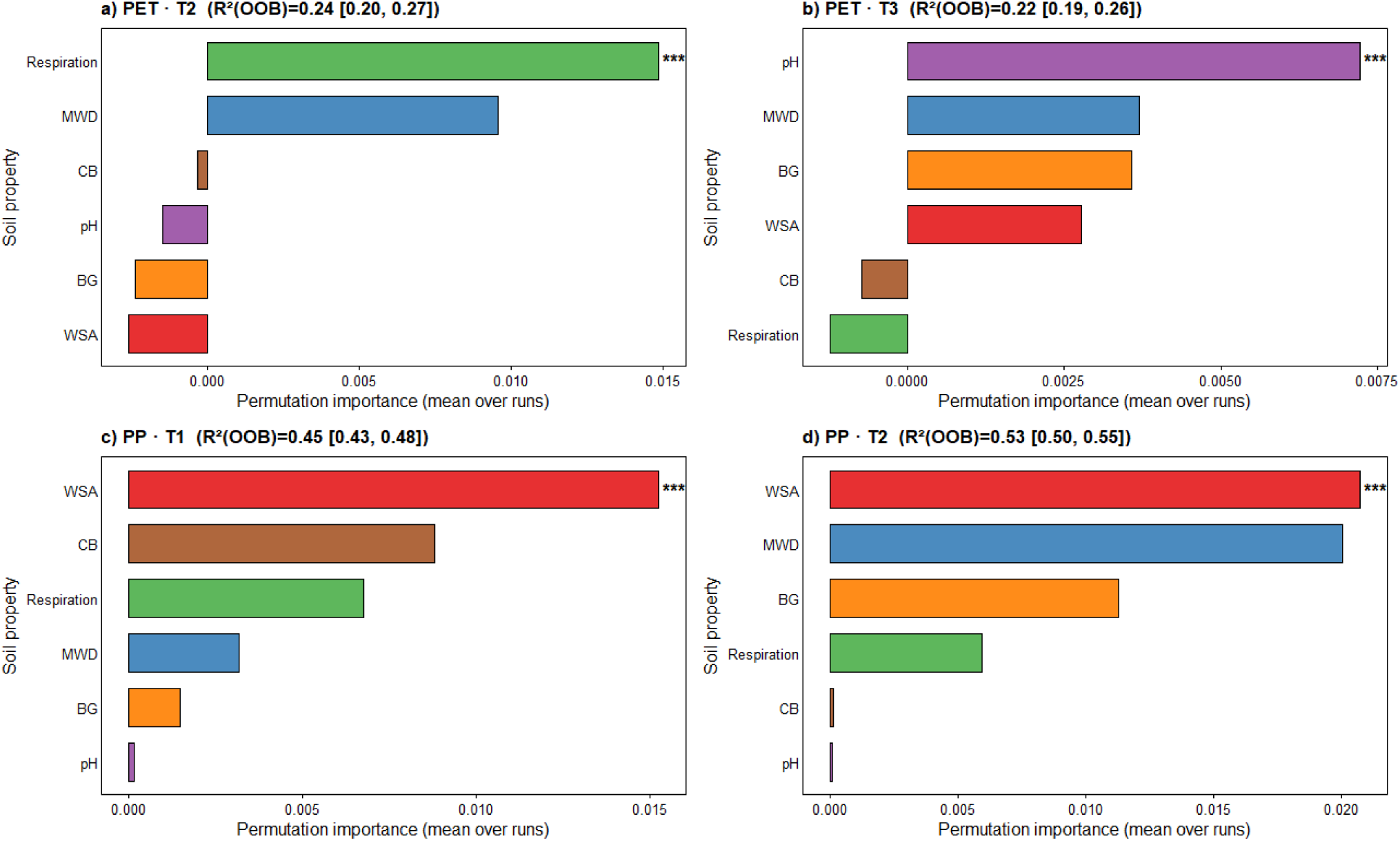
Random-forest variable importance analysis for soil multifunctionality prediction. PET: polyethylene terephthalate, PP: polypropylene. T1, T2 and T3 correspond to 2, 4, and 6 weeks of incubation, respectively. Panel a) PET-Timepoint 2; b) PET-Timepoint 3; c) PP-Timepoint 1; and d) PP-Timepoint 2. Bars represent relative importance of each variable for predicting soil function index based on the mean decrease in accuracy (absolute change in out-of-bag (OOB) error). R² values reflect OOB estimates, and empirical p-values of top variables were estimated via permutation analysis of predictor importance. Asterisks mark that all top-ranking-predictors importance was significantly greater than expected under random variation (empirical two-sided p_emp < 0.001 based on the permutation distribution of importance across runs). Several additional soil functions also showed significant importance, but only the dominant predictor was highlighted for simplicity to guide the subsequent mediation analysis. Soil aggregation metrics were consistently among the higher-ranking predictors overall, with WSA consistently ranking as the dominant predictor under PP.

Mediation analysis was performed to confirm whether microplastic dose influenced MFI indirectly through the ranked-top variables, and the results confirmed this mechanistic interpretation in one case. The PP concentration → WSA → MFI mediation was negative, with a 95% confidence interval excluding zero (indirect effect = −0.278 [−0.648, −0.037]; p = 0.010; R² = 0.46), indicating that reduced aggregate stability partially mediated the decline in multifunctionality after four weeks. In other words, MFI declined not simply because multiple functions changed independently, but because microplastics first destabilized soil structure, which subsequently affected other functions. However, when WSA was removed from the multifunctionality index structure, the indirect effect of PP concentration on MFI_no-WSA_ weakened, and its confidence interval overlapped zero.

Together, the multifunctionality analysis reveals a coherent ecological narrative: PP induces earlier, stronger system-level declines, with thresholds already emerging after two weeks of exposure; meanwhile, PET effects materialize later and require higher doses. Structural functions act as key mediators, determining whether soils retain the capacity to sustain multiple functions simultaneously. In short, these findings show that soil multifunctionality responds both gradually and abruptly to microplastic stress, with thresholds at 0.3% w/w PP after two weeks, and 0.55% w/w PET after six weeks, and that the reduction of soil physical structure is a relevant mechanism underlying declines in ecosystem-level performance.

## 4. Discussion and conclusions

### 4.1. Microplastics drive both gradual and abrupt responses across soil functions

Across all tested soil functions, most responses to increasing microplastic concentrations initially followed gradual and continuous trends rather than abrupt shifts, indicating that soil systems respond proportionally to microplastic inputs, and not with step change responses^63^. Nevertheless, after applying the significance filtering to retain only responses clearly driven by the concentration gradient, piecewise relationships became more prevalent, particularly at later incubation stages. This pattern indicates that cumulative physical disturbance may progressively weaken soil structure until a threshold that may exceed soil buffering capacity is reached^28,64^. However, profile-AIC uncertainty intervals were often wide, indicating limited precision in breakpoint estimates for individual functions. Therefore, reported breakpoints should be interpreted cautiously, as first approximations, and need to be further investigated in future studies.

Such abrupt shifts were more common on soil aggregate size, possibly due to alterations in soil porosity and aggregate stability. In some cases, these abrupt responses show interesting patterns in line with hormesis theory^33^. For instance, small additions of PP after six weeks did not significantly affect β-D-cellobiohydrolase, even suggesting a weak stimulation of this enzymatic activity. This may reflect a temporary enhancement of microbial activity due to an initial increase of carbon availability from the particles^21,25^. Once a critical concentration is exceeded, the response drops drastically, possibly associated with microbial stress or substrate limitation. This suggests that initial resource inputs associated with microplastic pollution may temporarily enhance or maintain microbial activity before structural degradation or particle toxicity overcomes functionality.

### 4.2. Microplastic-induced changes are polymer and time-specific

Responses of individual soil functions revealed clear polymer-specific patterns. For soil structural indicators, PP consistently reduced both WSA and MWD, particularly after four to six weeks of microplastic exposure, reflecting physical disturbance of aggregate size and stability. These effects are consistent with the particle morphology, as PP was present as small and rigid granular fragments. In contrast, PET generally produced weaker and more delayed structural effects, consistent with its smaller size as powder-like particles. These observations agree with previous studies^24,25^, where smaller, rigid, and angular particles, such as PP fragments, are more likely to lodge within pore spaces, interfering with particle binding forces. Meanwhile, softer or smoother PET powder particles may interact more weakly with soil structure, resulting in less severe structural changes.

Additionally, incubation time strongly modulated microplastic impacts. Early responses were predominantly gradual, suggesting that soil systems initially experienced limited disturbance. However, several properties exhibited abrupt transitions over time, and declines tended to intensify particularly under PP. These time-dependent patterns suggest that microplastics act through progressive physical restructuring. Initial aggregation, pore rearrangement, or microbial adaptation may temporarily buffer effects, but cumulative stress eventually leads to stronger losses on physical integrity and subsequent biochemical decline.

Overall, the factor ‘polymer type’ interacted strongly with incubation time. PET effects usually emerged later, possibly due to lower mechanical impact or slower interactions with microbial communities. In contrast, PP induced earlier functional changes, consistent with previous findings reporting faster biological and structural changes to this polymer type^65,66^. As mentioned before, this likely occurs because PP is often present as more rigid fragments with higher surface roughness. Together, these temporal trends indicate that exposure time is a key factor in assessing microplastic impacts on soil systems, and suggest that thresholds primarily arise after sufficient cumulative stress.

### 4.3. Microplastic-driven ecosystem thresholds

When individual soil properties were integrated into a multifunctionality index, distinct polymer and time-specific patterns appeared, as expected from previous evidence^24,25^. After six weeks of PET exposure, soil multifunctionality showed a negative slope trend at lower concentrations; however, after surpassing the estimated 0.55% w/w, soil multifunctionality seems to buffer and partially recover, suggesting a greater capacity to maintain coordinated functioning at higher concentrations. In contrast, PP exposure led to an earlier system-level decline around 0.3% w/w after two weeks, indicating lower resilience of soil multifunctionality to this polymer.

This low-dose system-level response, despite PP inducing thresholds at higher concentrations for individual properties, suggests that small structural changes across multiple functions can collectively amplify system-level responses. The disappearance of the breakpoint and its replacement by a linear decline after four weeks of PP exposure may indicate that the system moved beyond the threshold range, entering a state where multifunctionality decreases proportionally and potentially irreversibly with additional PP inputs. Differently, PET-induced changes appeared to remain buffered for longer, consistent with our GAM-based MWD results (Fig. 1a, 1b), where early declines may remain masked until a critical concentration is reached. These findings align with earlier studies reporting more rapid and pronounced effects of PP on soil functions^65,66^.

Machine-learning analyses reinforced the central role of soil structure in soil multifunctionality. Random forest models consistently captured a moderate amount of MFI variance under microplastic exposure (out-of-bag R^2^ = 0.22–0.53%), and identified soil aggregate stability as the dominant factor driving soil multifunctionality responses to PP pollution. Cheng et al.^18^ previously tested polyethylene and identified soil respiration as the main driver of soil multifunctionality, similar to our results for PET after four weeks of incubation (Fig 3.a.). However, they did not include soil physical measurements in their MFI analysis. In our study, the magnitude and significance of permutation importance indicated that aggregate stability was the strongest predictor of changes in multifunctionality (Fig. 3c, 3d), as previously reported, in a different context, by Eldridge et al.^67^. Thus, the primary importance of soil structure and its simultaneous control on multiple soil functions has previously been recognized^68,69^, yet here we specifically demonstrate that losses in aggregate stability under microplastic stress can predict declines in soil multifunctionality.

This relationship was independently confirmed by mediation analysis, which helped to provide a mechanistic insight. The pathway [MP] → WSA → MFI was significant for soils exposed to PP for four weeks, demonstrating that the decline in multifunctionality under microplastics is transmitted through reductions in soil aggregate stability rather than through direct biochemical shifts. Still, as the indirect effect turned non-significant after the robustness check, this should be interpreted cautiously, and further investigation is needed to explore this mechanistic explanation. However, this finding supports the overall linear decline of MFI under PP after four weeks of incubation. Overall, this shows that once soil aggregate stability is compromised and structural integrity is consistently lost, drastic shifts in biological and physicochemical performance follow, potentially leading to an earlier systemic response.

### 4.4. Synthesis and ecological implications

Taken together, our results converge on a coherent mechanistic explanation of multifunctionality decline. Soil ecosystems appear relatively resilient to microplastic inputs up to roughly 0.3–0.5% w/w, beyond which soils may still partially recover multifunctionality. However, after cumulative disturbance, physical integrity tends to be reduced, leading to cascading impacts in biological and physicochemical performance. This concentration window represents an ecologically meaningful transition from tolerance to degradation, conceptually equivalent to tipping points described in dryland or eutrophication systems^28,34^. Importantly, soil structural indicators consistently acted as early-warning indicators. The emergence of system-level thresholds in multifunctionality demonstrates that soil response to microplastic pollution cannot be understood through single functions: soil degradation arises from the coordinated decline of multiple processes, with soil structure change as the potential underlying mechanism.

To our knowledge, this study provides the first quantitative evidence of ecological thresholds in soil multifunctionality under microplastic pollution, establishing a mechanistic framework to further assess soil resilience at a systemic level. In short, our findings reveal that soil responses to increasing microplastic concentrations combine gradual trends with abrupt shifts for both individual functions and overall multifunctionality, depending on polymer type and time. Although exact threshold locations should be confirmed in future studies, the repeated emergence of breakpoints, particularly in soil structure and multifunctionality, highlights their ecological relevance. Overall, this estimated threshold range is ecologically critical to identify crucial concentrations under which soil systems transition from tolerance to degradation.

System-level performance consistently declined once concentrations reached approximately 0.3–0.5% w/w after two and six weeks of incubation, respectively, although some buffering capacity was occasionally observed. Machine-learning and mediation analyses converge on a common explanation: structural degradation, especially the loss of aggregate stability, seems to act as the primary driver linking microplastic inputs to declines in biological and chemical functioning. Together, these results pinpoint a critical microplastic concentration range indicating the transition from tolerance to degradation, providing novel quantitative insights into incipient ecological thresholds in soil multifunctionality under microplastic stress. Given projections suggesting that microplastic concentrations in agricultural soils may exceed 0.6% w/w under business-as-usual scenarios^11^, our findings have important implications for predicting long-term soil stability and for managing soil resilience in a world where microplastics are an increasingly widespread global change factor.

## Data availability statement

All data and code for analysis are publicly available on figshare: http://doi.org/10.6084/m9.figshare.31145929.

## Author contributions

T.M.-R: Conceptualization, Data curation, Formal analysis, Investigation, Visualization, Writing – original draft. M. Dacal: Conceptualization, Investigation, Writing – review & editing. E. Ring: Investigation, Writing – review & editing. M.C.-R: Conceptualization, Supervision, Writing – review & editing. All authors contributed to the final version of the manuscript and approved it for publication.

## Acknowledgments

This project has received funding from European Union’s HORIZON EUROPE research and innovation program GA N°101072777-PlasticUnderground HEUR-MSCA-2021-DN-01. MCR acknowledges additional support by the BMFTR-funded Rhizo4Bio project µPlastic (031B1410A). We acknowledge support by the Open Access Publication Initiative of Freie Universität Berlin.

