## Supplementary Materials for "Microplastics drive both linear and threshold-type shifts in soil multifunctionality along concentration gradients"

Tamara Meizoso-Regueira[1†], Marina Dacal[2,1†], Emma Ring[1†], Matthias C. Rillig[1,3*†]

1 Institute of Biology, Freie Universität Berlin, Altensteinstr. 6 14195 Berlin, Germany

2 Grupo de Ecoloxía Animal (GEA), Universidad de Vigo, 36310 Vigo, Spain

3 Berlin-Brandenburg Institute of Advanced Biodiversity Research (BBIB), Berlin, Germany

†**ORCID:** Tamara Meizoso-Regueira: https://orcid.org/0000-0003-0743-9836; Marina Dacal: https://orcid.org/0000-0002-1321-9373; Emma Ring: https://orcid.org/0009-0000-5587-9356; Matthias C. Rillig: https://orcid.org/0000-0003-3541-7853

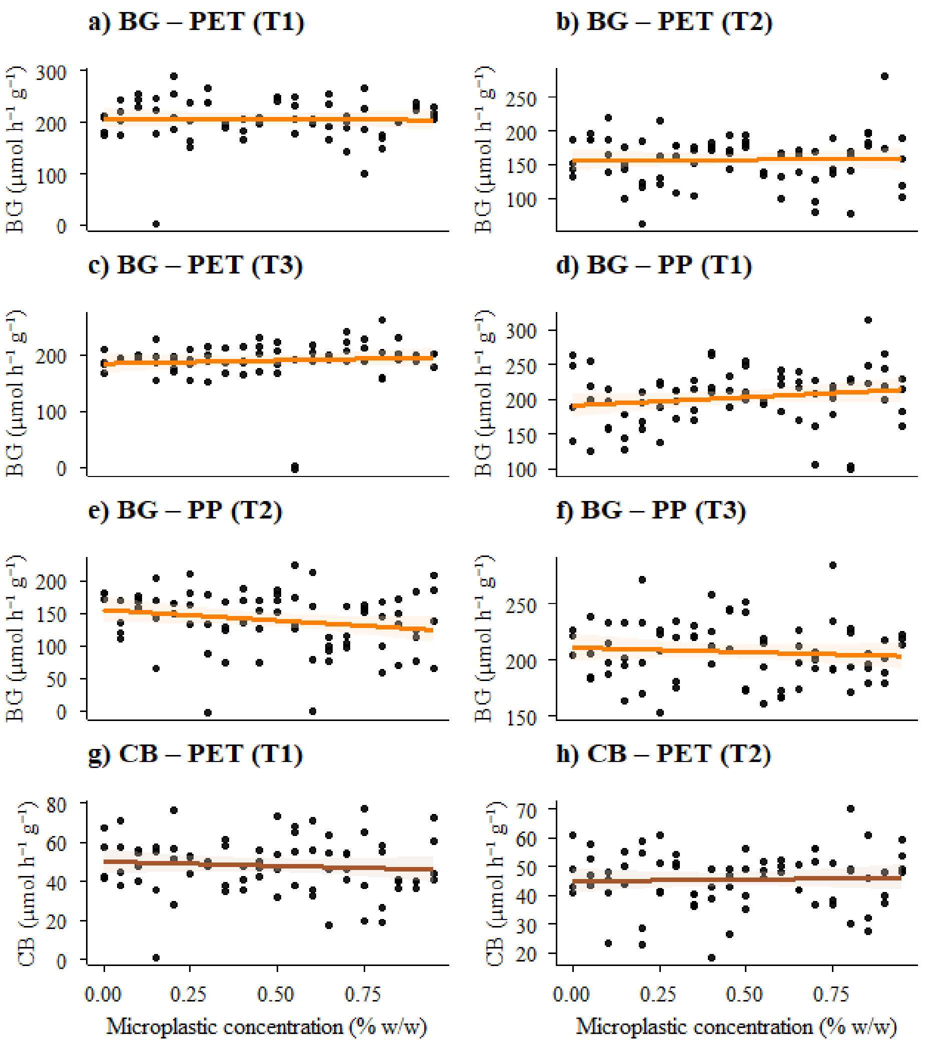


**Figure S1. Winner models by AIC comparison that did not pass the significance criteria for β-D-glucosidase (BG) and cellobiohydrolase (CB) activity**. PET: polyethylene terephthalate, PP: polypropylene. T1, T2 and T3 correspond to 2, 4, and 6 weeks of incubation, respectively. Linear and quadratic models were tested using the overall F-test (a stricter criterion), while GAM and piecewise models lack an equivalent parametric F-test and were instead evaluated via smooth or slope term significance. Consequently, most discarded winners corresponded to linear fits.

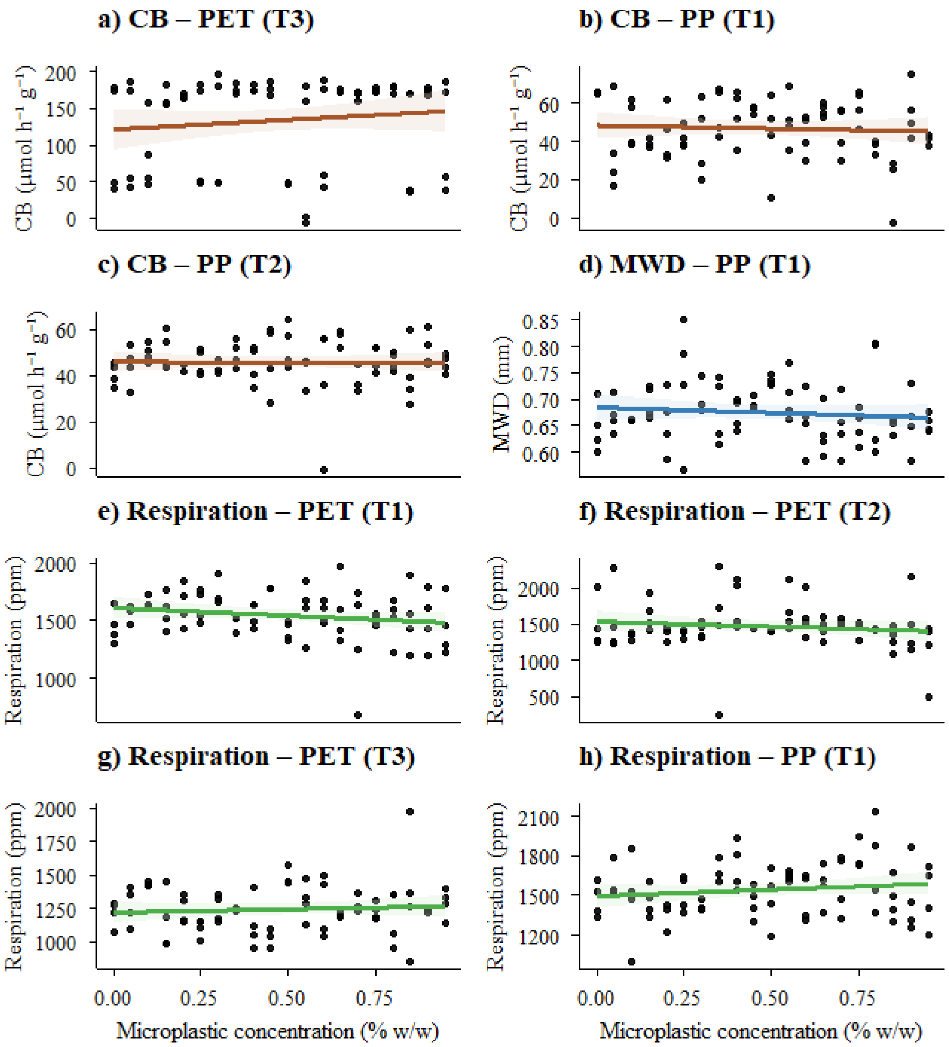


**Figure S2. Winner models by AIC comparison that did not pass the significance criteria for β-D-cellobiohydrolase (CB), mean-weight diameter (MWD) and respiration functions.** PET: polyethylene terephthalate, PP: polypropylene. T1, T2 and T3 correspond to 2, 4, and 6 weeks of incubation, respectively. Linear and quadratic models were tested using the overall F-test (a stricter criterion), while GAM and piecewise models lack an equivalent parametric F-test and were instead evaluated via smooth or slope term significance. Consequently, most discarded winners corresponded to linear fits.

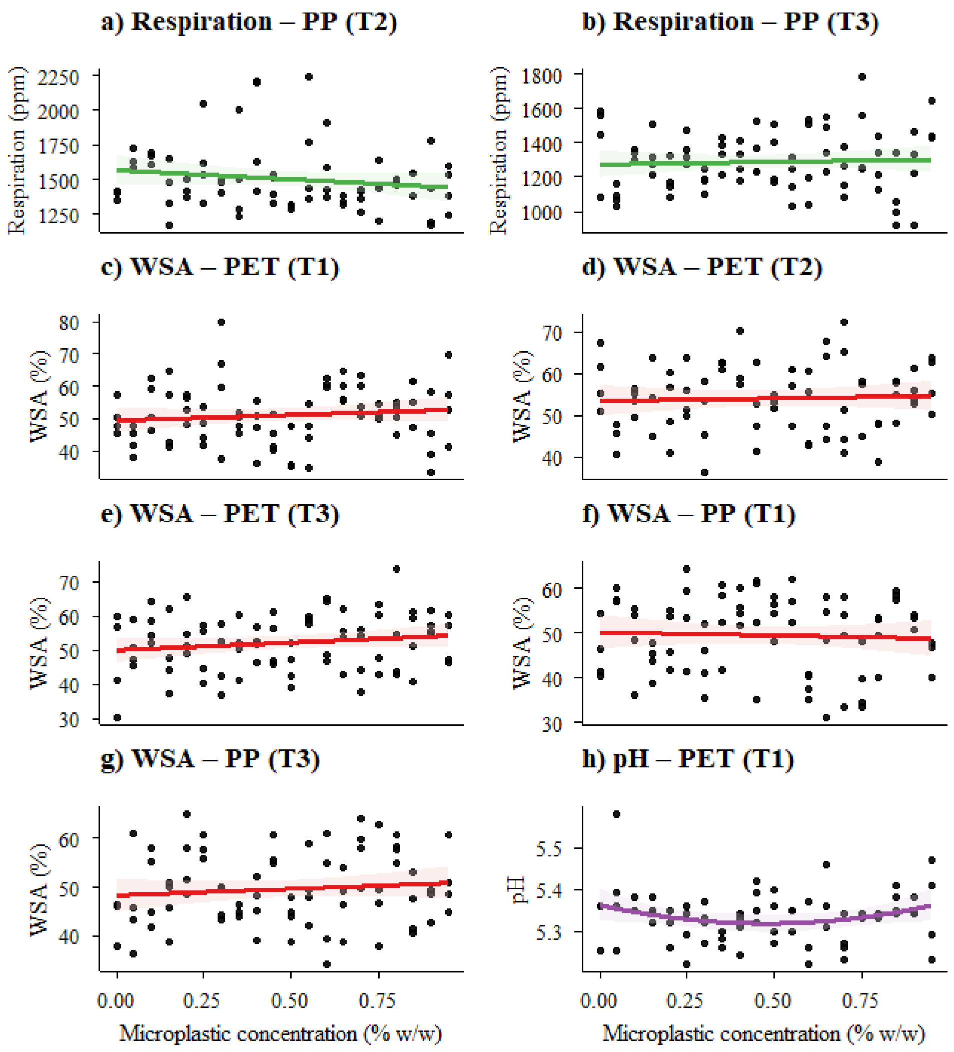


**Figure S3. Winner models by AIC comparison that did not pass the significance criteria for respiration, water-stable aggregates (WSA) and pH functions.** PET: polyethylene terephthalate, PP: polypropylene. T1, T2 and T3 correspond to 2, 4, and 6 weeks of incubation, respectively. Linear and quadratic models were tested using the overall F-test (a stricter criterion), while GAM and piecewise models lack an equivalent parametric F-test and were instead evaluated via smooth or slope term significance. Consequently, most discarded winners corresponded to linear fits.

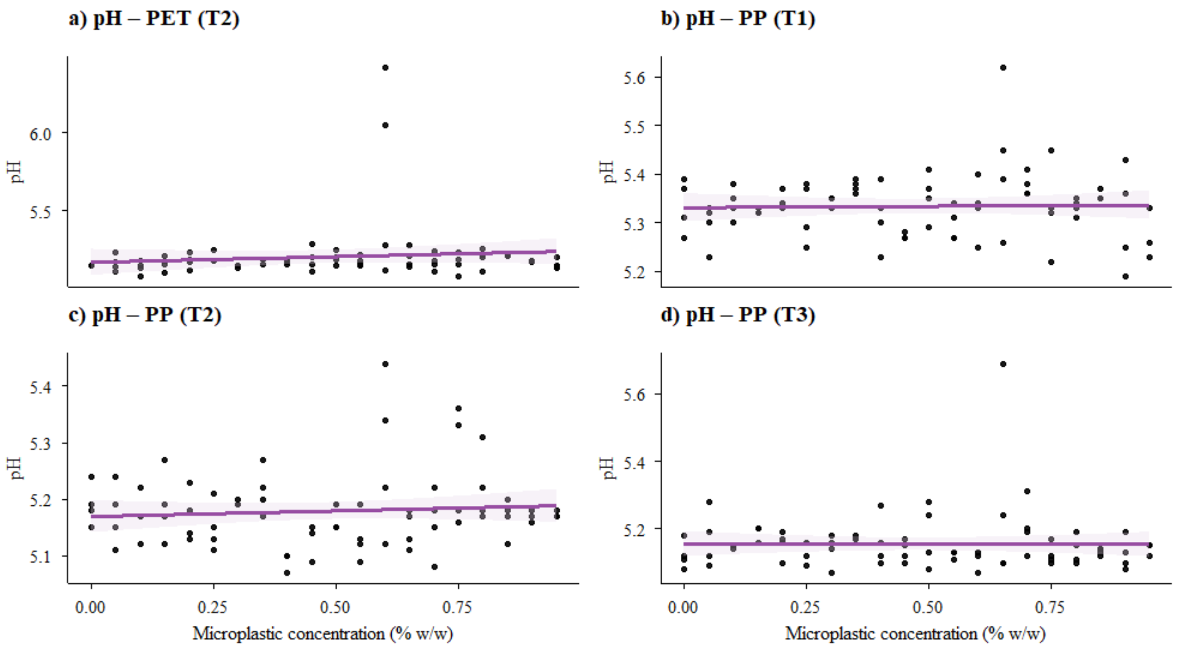


**Figure S4. Winner models by AIC comparison that did not pass the significance criteria for pH function.** PET: polyethylene terephthalate, PP: polypropylene. T1, T2 and T3 correspond to 2, 4, and 6 weeks of incubation, respectively. Linear and quadratic models were tested using the overall F-test (a stricter criterion), while GAM and piecewise models lack an equivalent parametric F-test and were instead evaluated via smooth or slope term significance. Consequently, most discarded winners corresponded to linear fits.

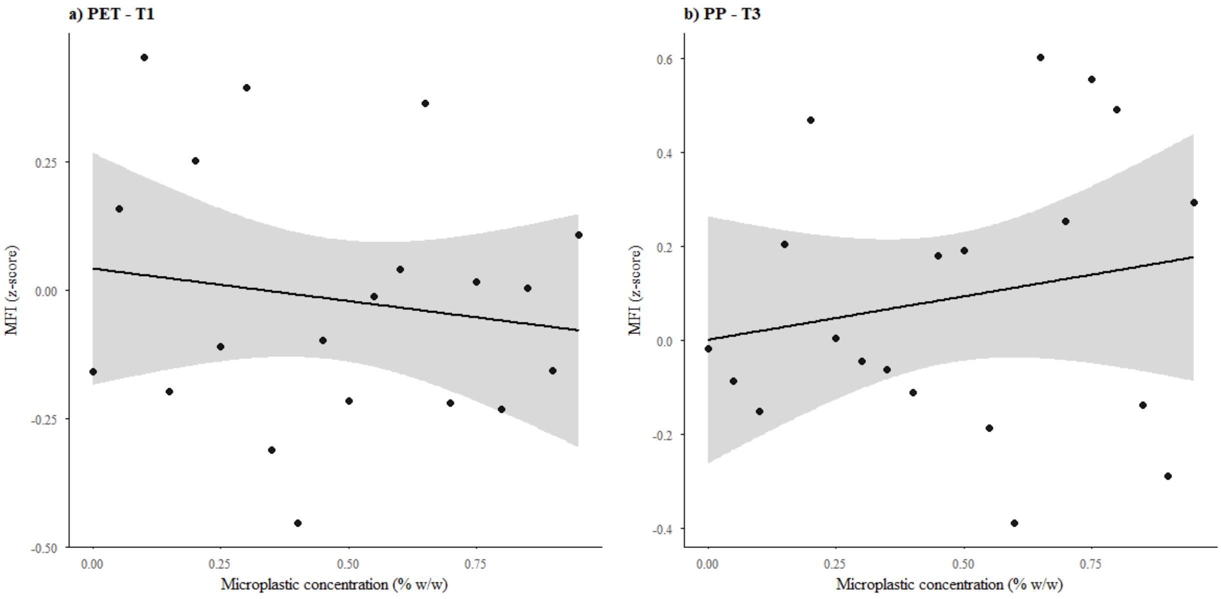


**Figure S5. Winner models by AIC comparison for soil multifunctionality index (MFI) that did not pass the significance criteria.** PET: polyethylene terephthalate, PP: polypropylene. T1 and T3 correspond to 2 and 6 weeks of incubation, respectively. From top to bottom: a) MFI corresponding to PET-Timepoint 1 (linear); b) PP-Timepoint 3 (linear).

**Table S1. Goodness-of-fit metrics to evaluate and compare non-significant model fits for each combination of polymer x timepoint for β-D-glucosidase (BG) and cellobiohydrolase (CB), mean weight diameter (MWD) and respiration (RES), including AIC, BIC and $R^2$.** Highlighted in bold are the winner models following our $\Delta AIC \geq 2$ criteria, prioritizing parsimony (l=linear, nl=nonlinear, p=piecewise). The term p-val corresponds to the overall F-test for linear and quadratic models, or to the significance of the smooth and slope/step terms for GAM and piecewise models, respectively. We do not report the estimated breakpoints $\hat{\psi}$ along with profile-AIC 95% confidence intervals as linear models (only directly compared against GAM or quadratic) consistently presented the best fit, so there was no threshold to be reported.

| Property | $AIC_l$ | $AIC_{nl}$ | $AIC_p$ | $BIC_l$ | $BIC_{nl}$ | $BIC_p$ | $R^2_l$ | $R^2_t$ | $\hat{\psi}$ (CIs) | p-val |
|---|---|---|---|---|---|---|---|---|---|---|
| **BG-PET-T1** | **724.6** | 731.4 | NA | 724.6 | 731.4 | NA | 0 | NA | NA | 0.861 |
| **BG-PET-T2** | **746.7** | 753.6 | NA | 746.7 | 753.6 | NA | 0 | NA | NA | 0.932 |
| **BG-PET-T3** | **762.1** | 769 | NA | 762.4 | 770.5 | NA | 0.007 | NA | NA | 0.467 |
| **BG-PP-T1** | **790.6** | 797.6 | NA | 790.6 | 797.6 | NA | 0.029 | NA | NA | 0.139 |
| **BG-PP-T2** | **808.6** | 815.6 | NA | 808.6 | 815.6 | NA | 0.04 | NA | NA | 0.081 |
| **BG-PP-T3** | **709.3** | 716.3 | NA | 709.3 | 716.3 | NA | 0.007 | NA | NA | 0.472 |
| **CB-PET-T1** | **615.7** | 622.7 | NA | 615.7 | 622.7 | NA | 0.007 | NA | NA | 0.465 |
| **CB-PET-T2** | **551.8** | 558.7 | NA | 552.1 | 560 | NA | 0.001 | NA | NA | 0.753 |
| **CB-PET-T3** | **780.8** | 787.6 | NA | 780.8 | 787.6 | NA | 0.016 | NA | NA | 0.299 |
| **CB-PP-T1** | **617.1** | 624 | NA | 617.1 | 624 | NA | 0.004 | NA | NA | 0.61 |
| **CB-PP-T2** | **557.8** | 564.8 | NA | 557.8 | 564.8 | NA | 0 | NA | NA | 0.95 |
| **MWD-PP-T1** | **-222.4** | -215.4 | NA | -224.2 | -214.8 | NA | 0.011 | NA | NA | 0.361 |
| **RES-PET-T1** | **1013.3** | 1020.2 | NA | 1013.7 | 1022.1 | NA | 0.036 | NA | NA | 0.104 |
| **RES-PET-T2** | **1053.3** | 1060.1 | NA | 1052.8 | 1062 | NA | 0.017 | NA | NA | 0.271 |
| **RES-PET-T3** | **965.2** | 972 | NA | 965.2 | 972 | NA | 0.007 | NA | NA | 0.491 |

**Table S2. Goodness-of-fit metrics to evaluate and compare non-significant model fits for each combination of polymer x timepoint for respiration (RES), water-stable aggregates (WSA) and pH, including AIC, BIC and $R^2$.** Highlighted in bold are the winner models following our $\Delta AIC \geq 2$ criteria, prioritizing parsimony (l=linear, nl=nonlinear, p=piecewise). The term p-val corresponds to the overall F-test for linear and quadratic models, or to the significance of the smooth and slope/step terms for GAM and piecewise models, respectively. We do not report the estimated breakpoints $\widehat{\psi}$ along with profile-AIC 95% confidence intervals as linear models (only directly compared against GAM or quadratic) consistently presented the best fit, so there was no threshold to be reported.

| Property | $AIC_l$ | $AIC_{nl}$ | $AIC_p$ | $BIC_l$ | $BIC_{nl}$ | $BIC_p$ | $R^2_l$ | $R^2_t$ | $\widehat{\psi}$ (CIs) | p-val |
|---|---|---|---|---|---|---|---|---|---|---|
| **RES-PP-T1** | **1013.2** | 1013.6 | NA | 1020.2 | 1021.9 | NA | 1020.2 | NA | NA | 0.219 |
| **RES-PP-T2** | **990.8** | 990.8 | NA | 997.6 | 997.7 | NA | 997.6 | NA | NA | 0.18 |
| **RES-PP-T3** | **1030.5** | 1030.5 | NA | 1037.6 | 1037.6 | NA | 1037.6 | NA | NA | 0.669 |
| **WSA-PET-T1** | **573.2** | 573.2 | NA | 580.3 | 580.3 | NA | 580.3 | NA | NA | 0.322 |
| **WSA-PET-T2** | **521.5** | 521.5 | NA | 528.4 | 528.4 | NA | 528.4 | NA | NA | 0.698 |
| **WSA-PET-T3** | **561.8** | 561.8 | NA | 568.9 | 568.9 | NA | 568.9 | NA | NA | 0.174 |
| **WSA-PP-T1** | **538.8** | 538.8 | NA | 545.7 | 545.7 | NA | 545.7 | NA | NA | 0.672 |
| **WSA-PP-T3** | **519** | 519 | NA | 526 | 526 | NA | 526 | NA | NA | 0.366 |
| **pH-PET-T1** | **-191** | -193.1 | NA | -184.2 | -184 | NA | -184.2 | NA | NA | 0.985 |
| **pH-PET-T2** | **-37.2** | -37.9 | NA | -30.3 | -28.8 | NA | -30.3 | NA | NA | 0.335 |
| **pH-PP-T1** | **-191.9** | -191.8 | NA | -185 | -182.6 | NA | -185 | NA | NA | 0.829 |
| **pH-PP-T2** | **-194.2** | -194.2 | NA | -187.2 | -187.2 | NA | -187.2 | NA | NA | 0.443 |
| **pH-PP-T3** | **-157.8** | -157.8 | NA | -150.9 | -150.8 | NA | -150.9 | NA | NA | 0.961 |

**Table S3. Goodness-of-fit metrics to evaluate and compare non-significant model fits for each combination of MFI x polymer x timepoint, including AIC, BIC and $R^2$.** Highlighted in bold are the winner models following our $\Delta AIC \geq 2$ criteria, prioritizing parsimony (l=linear, nl=nonlinear, p=piecewise). The term p-val corresponds to the overall F-test for linear and quadratic models, or to the significance of the smooth and slope/step terms for GAM and piecewise models, respectively. We do not report the estimated breakpoints $\hat{\psi}$ along with profile-AIC 95% confidence intervals as linear models (only directly compared against gam or quadratic) consistently presented the best fit, so there was no threshold to be reported.

| **Property** | **$AIC_l$** | **$AIC_{nl}$** | **$AIC_p$** | **$BIC_l$** | **$BIC_{nl}$** | **$BIC_p$** | **$R^2_l$** | **$R^2_t$** | **$\hat{\psi}$** | **p-val** |
|---|---|---|---|---|---|---|---|---|---|---|
| **MFI-PET-T1** | **5.2** | 5.2 | NA | 8.2 | 8.2 | NA | 0.023 | NA | NA | 0.52 |
| **MFI-PP-T3** | **11.2** | 11.2 | NA | 14.2 | 14.2 | NA | 0.036 | NA | NA | 0.42 |
